# Hyperdisordered cell packing on a growing surface

**DOI:** 10.1101/2024.05.20.593453

**Authors:** R. J. H. Ross, Giovanni D. Masucci, Chun Yen Lin, Teresa L. Iglesias, Sam Reiter, Simone Pigolotti

## Abstract

While the physics of disordered packing in non-growing systems is well understood, unexplored phenomena can emerge when packing takes place in growing domains. We study the arrangements of pigment cells (chromatophores) on squid skin as a biological example of a packed system on an expanding surface. We find that relative density fluctuations in cell numbers grow with spatial scale. We term this behavior “hyperdisordered”, in contrast with hyperuniform behavior in which relative fluctuations tend to zero at large scale. We find that hyperdisordered scaling, akin to that of a critical system, is quantitatively reproduced by a model in which hard disks are randomly inserted in a homogeneously growing surface. In addition, we find that chromatophores increase in size during animal development, but maintain a stationary size distribution. The physical mechanisms described in our work may apply to a broad class of growing dense systems.

## I. INTRODUCTION

Many physical and biological systems are constituted by dense, disordered arrangements of individual units. Examples include liquids [1], glasses [2], granular systems [3], packed macroscopic objects [4–6], bacterial populations [7], and multicellular tissues [8–11]. The study of these systems has led to the discovery of broad physical properties that characterize dense disordered packing.

One such property is hyperuniformity. To define it, we consider how the average number ⟨*N*⟩ of units and their variance 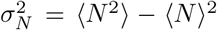 scale at increasing sample areas in a dense homogeneous system. For large areas, one expects a relation of the form

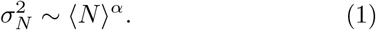

If units were independently placed at random, their number in each area would follow a Poisson distribution, implying *α* = 1. Hyperuniform systems are characterized by having *α <* 1 [12]. This means that they exhibit relative density fluctuations that are suppressed at large spatial scales [12–14]. Originally studied in maximally random jammed systems [6, 12], hyperuniformity has subsequently been observed in several models in non-equilibrium statistical physics [15–17] and in active systems [18]. Hyperuniformity has also been observed in biological systems, such as in the arrangement of cells in the avian retina [19] and in leaf vein networks [20].

Growth is a prominent aspect of living tissues, which sets them apart from prototypical examples of dense disordered systems in physics. The consequences of growth for cell arrangements have been studied in a limited way so far. Modeling shows that growth can control pattern formation and alter cell population dynamics [21–25]. It is thus conceivable that the presence of growth can lead to new physics in dense disordered systems. Experimentally, testing of these ideas has been limited, largely due to the difficulty in quantitatively and precisely measuring tissue arrangements during growth.

In this paper, we assess the impact of tissue growth on a dense arrangement of cells. Our experimental model system is the arrangement of pigment cells (chromatophores) on the skin of the oval squid. Combining theory and experiments, we reveal two striking properties of chromatophore patterns. First, we find that they are “hyperdisordered”, i.e., characterized by *α >* 1. This atypical behavior can be seen as the opposite of hyperuniform systems, and generically emerges as a consequence of packing and growth. Second, the size distribution of chromatophores is stationary during growth. Our theory shows that this stationary distribution is only achievable via an aging mechanism, whereby individual cells possess some notion of the squid age. This prediction is in excellent quantitative agreement with our experimental measurements. Together, our results reveal fundamental physical mechanisms governing the dynamics of dense growing physical systems.

## II. HYPERDISORDERED SCALING OF CHROMATOPHORE POINT PATTERNS

### A. Experimental system

Throughout development, the skin of the oval squid is populated by pigment-filled cells called chromatophores. Insertion of a chromatophore is thought to be possible only if the new chromatophore is at a minimum exclusion distance from preexisting ones [26, 27]. Following insertion into the skin, chromatophores do not move. This means that squid chromatophores, besides being the constitutive elements of a point pattern, can also function as reference points to precisely determine skin growth. To perform functions related to camouflage and communication, chromatophores can be expanded beyond their resting size through neurally controlled contraction of muscles surrounding each chromatophore [28]. How-ever, throughout this work we used anaesthetised animals to focus exclusively on the resting size. A chromatophore’s resting size (which we simply refer to as “chromatophore size” from now on) tends to increase as the chromatophore ages [29].

To study chromatophore patterns, we took highresolution images of the mantles of 10 different squids of the same age, see Fig. 1a and 1b. The experimental protocol and imaging procedure are detailed in Appendix A.

**FIG. 1.**
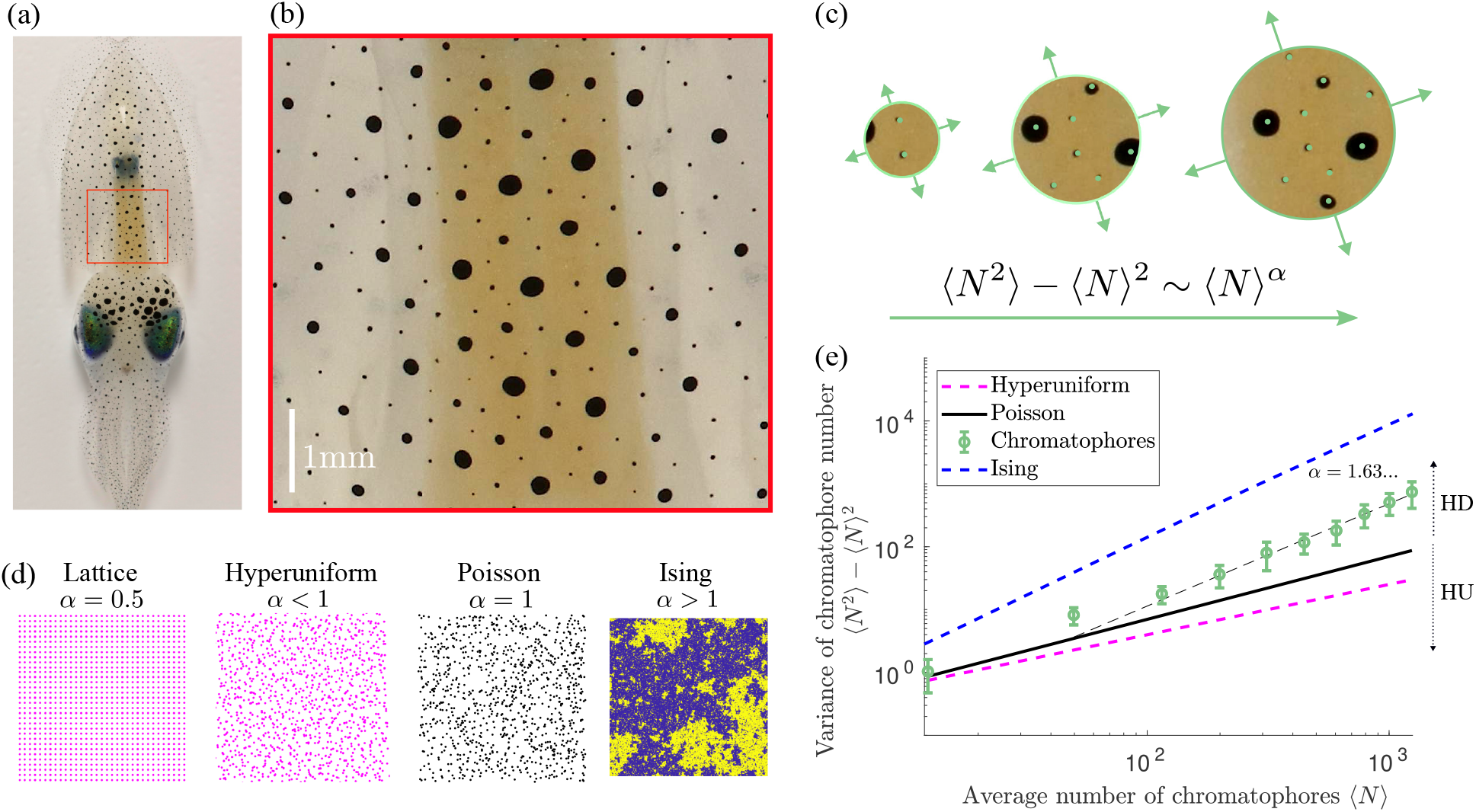
Chromatophore patterns are hyperdisordered. (a) An approximately 6-weeks old squid, viewed from the top. (b) Magnification of the mantle region indicated by the red box in (a). (c) Number of chromatophores *N* sampled in circular areas of increasing radius *R*. (d) Examples of (left to right) a uniform pattern, a hyperuniform pattern, a Poisson pattern, and a configuration of the two-dimensional Ising model at the critical temperature. (e) Chromatophore number variance, as a function of chromatophore number within an expanding area for the synthetic patterns in (d), and for squid chromatophores, which show hyperdisordered scaling (HD) (Total number of chromatophores *N*_tot_ = 17140 aggregated from 10 squids). Here and in the following, error bars denote standard deviations, unless stated otherwise.

### B. Chromatophore scaling

Our images of squid mantles show that chromatophores are arranged in a point pattern, characterized by larger, older chromatophores being surrounded by smaller, younger ones, see Fig. 1a and 1b. Visually, chromatophores appear at a characteristic distance from each other, forming a lattice-like structure. The density of chromatophores in our imaging area appears to be uniform, see Appendix B.

To characterize the degree of ordering of chromatophore patterns, we measure how the chromatophore number variance 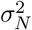 scales with the average number ⟨*N*⟩ of chromatophores at increasing sample areas (Fig. 1c) [12]. Since chromatophore patterns (Fig. 1b) emerge through packed insertion into the skin, one may expect them to exhibit hyperuniform scaling, see Fig. 1d. However, since the system is growing, small gaps are constantly being created between chromatophores, resulting in cell arrangements that substantially differ from those of commonly studied jammed systems. In fact, we observe that the number variance grows super-Poissonianly, i.e., *α >* 1, see Fig. 1e. We term this behavior “hyperdisordered”. Hyperdisordered systems share some properties with critical physical systems, such as large fluctuations at large scales (Fig. 1d).

### C. Squid model: hard disks on an expanding surface

To rationalize the observed hyperdisordered behavior, we introduce a model in which chromatophores are inserted as the mantle grows. This growth is linear in length and uniform. A chromatophore insertion is permitted only if the distance between the candidate new chromatophore and all the existing ones is larger than an exclusion distance Δ, see Fig. 2a. In this respect, the model can be thought of as a system of randomly placed hard disks on a homogeneously growing surface. The linear growth rate and exclusion distance are determined by matching the surface growth rate and spatial density of chromatophores from experiments. Additional details on the model simulations and parameter choices are presented in Appendix C. Despite its simplicity, the model generates patterns presenting hyperdisordered scaling of the number variance, consistent with that observed in real squids (Fig. 2b).

**FIG. 2.**
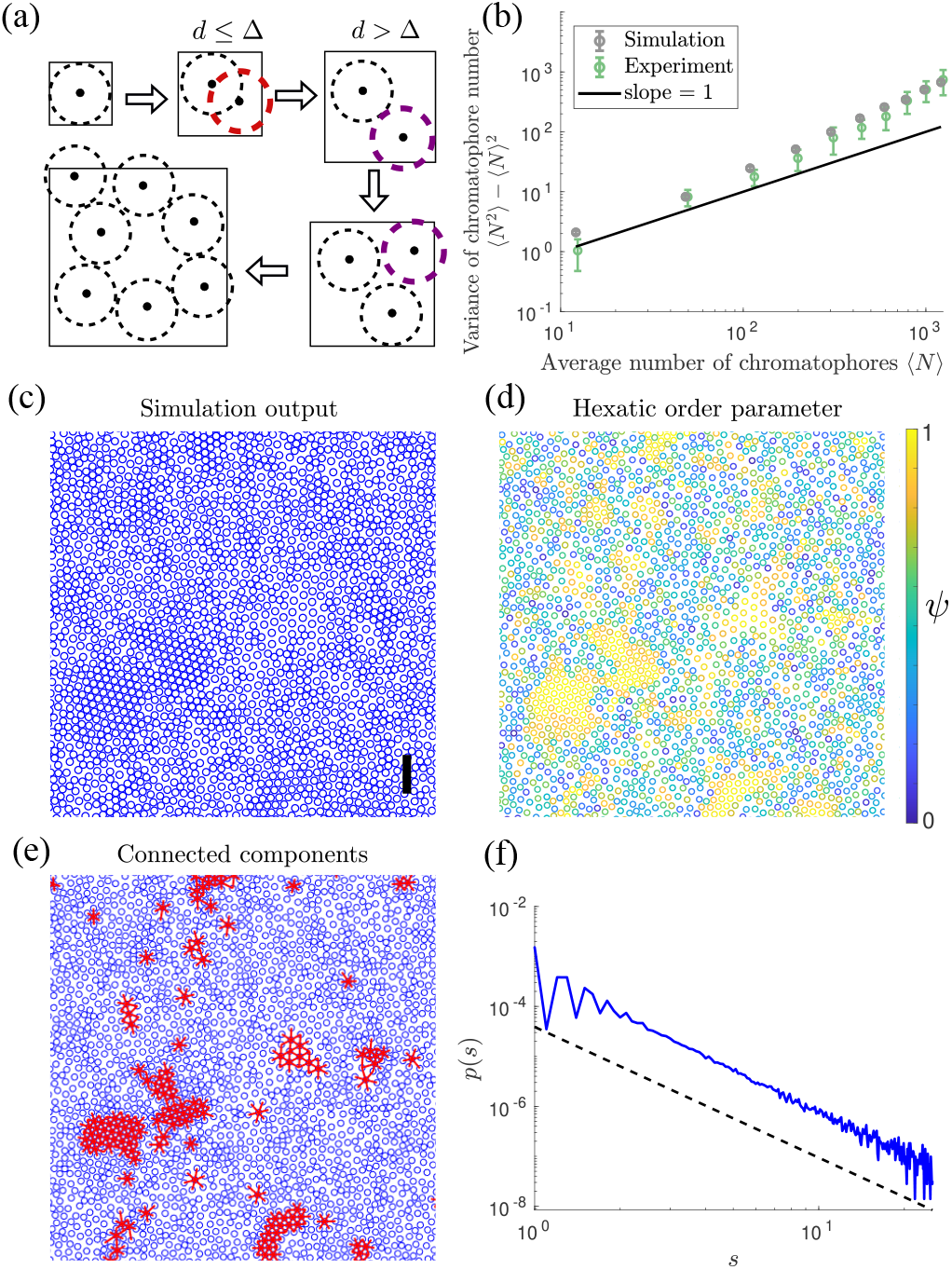
Growth generates hyperdisordered scaling. (a) Schematic of the model. Black dots represent chromatophores and dashed circles their exclusion areas, which are identical for all chromatophores. The red dashed circle represents an event in which an attempted chromatophore insertion is rejected because the distance between chromatophore centres is too small. The purple dashed circle represents a successful chromatophore insertion. (b) Number variance scaling (system size of *N* = 10^4^ chromatophores averaged over 500 simulations). (c) A configuration of the squid model (see SM Movie 1). The black scale bar indicates 1mm; the simulation area is 10mm^2^. (d) Hexatic order parameter *ψ* for each cell in (c), see Eq. (2). (e) Red domains represent the connected components of the top 5% cells in (d), ordered according to their value of |*ψ*_*j*_|. (f) Size distribution of the red domains shown in (e). The black-dashed line represents a power law with exponent -2.62.

We next seek to understand the cause of this scaling behavior. A large-scale simulation of this model shows nearly regularly packed domains, with different densities and without a clear characteristic size, surrounded by more disordered regions, see Fig. 2c. During growth, the density of these domains oscillates in a sawtooth manner. The reason for this is that the domains gradually become less packed as the surface grows, until there is enough space for inserting new chromatophores in the gaps between existing ones, see SM Movie 1. To quantify the statistics of these domains, we introduce the hexatic order parameter

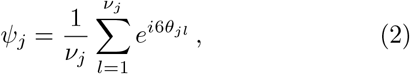

where *j* is an index representing a given cell, *v*_*j*_ is the number of nearest neighbours of cell *j* computed via Delauney triangulation, and *θ*_*jl*_ is the angle between the vector pointing from cell *j* to *l* and the *x* axis. A large value of |*ψ*_*i*_| can be interpreted as a local ordering of the neighbors of cell *j*, see Fig. 2d. We therefore identify domains as connected sets of neighboring cells characterized by high values of |*ψ*_*j*_|, see Fig. 2e. We find that the size distribution of these domains is well described by a power law (Fig. 2f), consistent with the idea that the dynamical heterogeneity generated by the interplay of growth and area exclusion is scale free.

Hyperdisordered scaling of a point pattern is necessarily associated with a pair correlation function whose integral in space diverges [6]. The envelope of the correlation function in the squid and in the model appear to decay in a similar way, see Fig. 3a. This decay is compatible with *r*^*−*2^, so that the integral in space of the correlation function presents a logarithmic divergence in two dimensions. This behavior is more evident in a large-scale simulation of the model, see Fig. 3b.

**FIG. 3.**
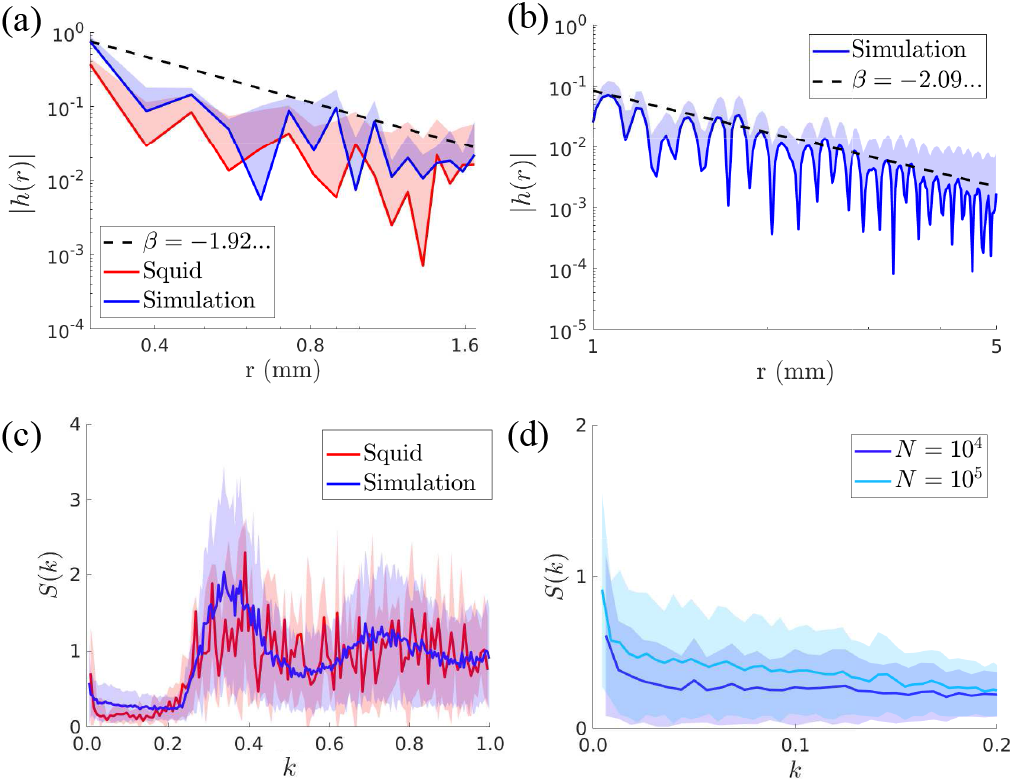
Correlation functions and structure factors are consistent with hyperdisordered behavior. (a) Comparison of correlation functions from experiment and simulation, for comparable numbers of data points. The correlation function is defined as *h*(*r*) = *ρ*_2_(*r*)*/ρ*^2^ *−*1, where *ρ*_2_(*r*) is the density of pairs at distance *r* and *ρ* is the one-body density. The black dashed line is a fit of the envelope of the simulation data. (b) Correlation function in the squid model for large system size. (c) Comparison of structure factor for squid chromatophores and from simulation data. (system size of *N* = 10^4^ chromatophores averaged over 500 simulations) (d) Comparison of structure factors from the squid model for different system sizes.

Another equivalent property of a hyperdisordered system is that the structure factor

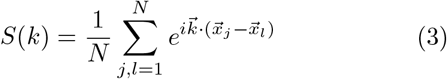

must diverge in the limit *k→* 0. However, this property is hard to test in practice, as taking this limit requires a very large system. The structure factor measured in the squid presents an increasing behavior as *k→* 0, see Fig. 3c. A similar behavior is observed in the model. Running simulations of the model for larger system size reveals a more pronounced peak of the structure factor near the origin, see Fig. 3d, as expected for a hyperdisordered system.

### D. One-dimensional iterative model

Why does growth lead to hyperdisordered behavior in a dense system? To gain understanding, we propose a one-dimensional iterative model that recapitulates some of the salient features of our system. In this model, a 1D domain is initialised with *n* cells placed uniformly at random. In analogy with the squid model, we call Δ the typical distance between cells. We set Δ = 1, so that an initial domain of integer length *L*, contains *n* = *L* initial cells. At each step of the model (generation), the domain size is doubled and new cells are placed at *x*_*i*_ + 𝒩 (Δ, *γ*), where *x*_*i*_ are the positions of the cells already present and 𝒩 is a Gaussian random variable with mean Δ and variance *γ* (Fig. 4a). This process is repeated for *t* steps, so that the final domain size is *n ×*2^*t*^.

**FIG. 4.**
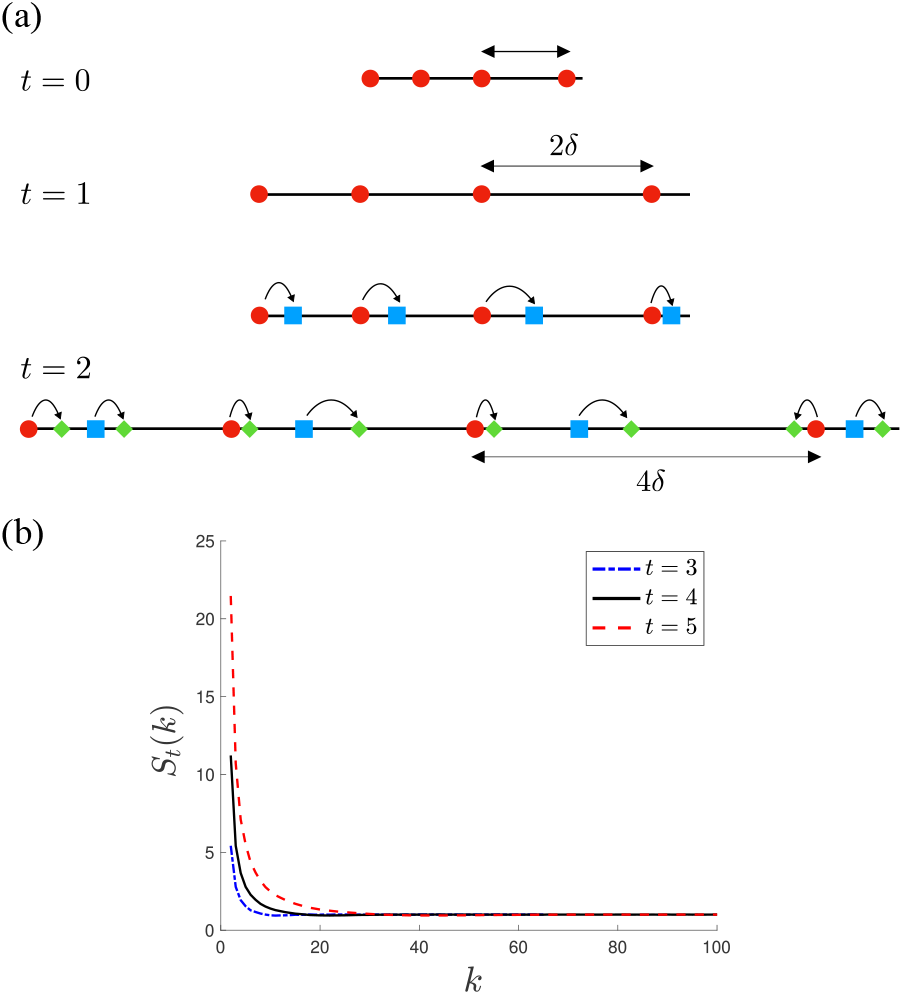
The iterative model presents hyperdisordered behavior. (a) Schematic of the model. (b) Structure factor as expressed by Eq. (4) for *L* = 4, *γ* = 1, and *t* = 3, 4, 5 as shown in the legend.

To analyze the model, we define the structure factor *S*_*t*_(*k*) of the model at generation *t*. We recall that a point pattern is hyperdisordered if the structure factor diverges as *k→* 0. We find that the structure factor is expressed by

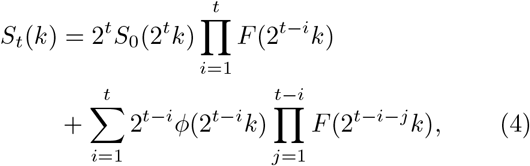

where

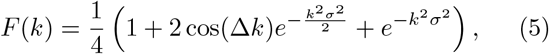

and

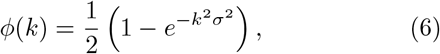

see Appendix D for a derivation. In the limit of large *t* and *k→* 0, the structure factor given in Eq. (4) diverges (see Appendix D and Fig. 4b), implying that the behavior of the iterative model is hyperdisordered, regardless of parameter choice. This suggests that, in essence, hyperdisordered behavior is caused by growth exporting density fluctuations to increasingly large scales.

## III. DYNAMICS OF CHROMATOPHORE SIZE PRESENTS AGING

### A. Stationary distribution of chromatophore size

We now study the dynamical behavior of chromatophore patterns, in particular how chromatophore sizes evolve as the system grows. To track populations of chromatophores on growing patches of squid skin through time, we employed three-dimensional imaging to construct the skin manifold (Fig. 5a), and used computer vision techniques to segment chromatophores and locate their center of mass, see Fig. 5b and Appendix A. Tracking an initial set of chromatophores over 6 weeks reveals uniform linear growth in the inter-chromatophore distance over time, and consequently also in the system size (Fig. 5c). In parallel, individual chromatophores grow in size as they age (Fig. 5b). Increases in chromatophore radii and separation are offset by the insertion of small new chromatophores into the skin, resulting in an approximately constant chromatophore density and stationary size distribution over time, see Fig. 5d and 5e. The stationarity of the chromatophore size distribution is confirmed by a Kolmogorov-Smirnoff test, see Appendix E.

**FIG. 5.**
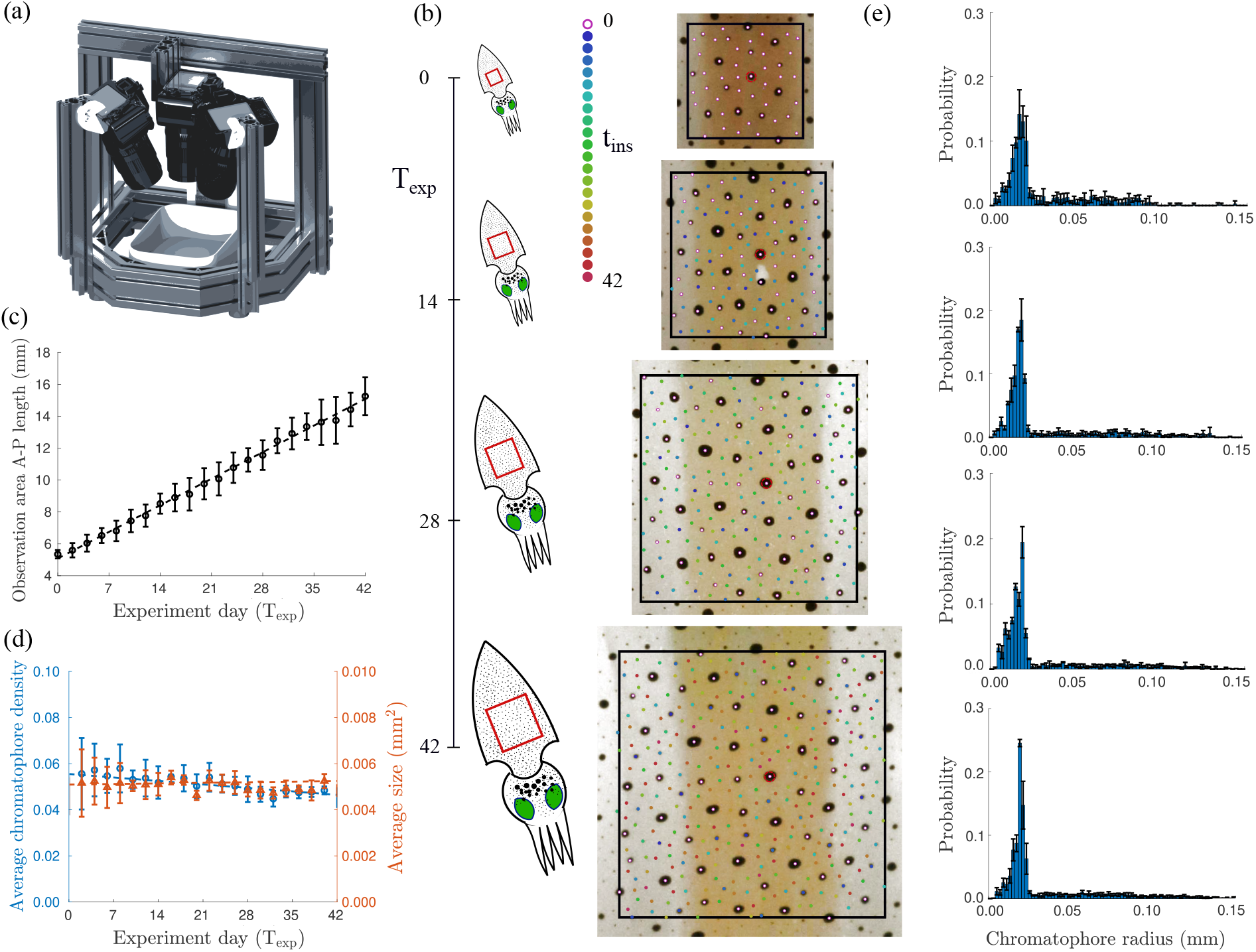
Chromatophore size distribution is stationary during growth. (a) Experimental setup for 3D imaging of squid skin. (b) Left: schematic of chromatophore tracking experiment. Right: skin patch with tracked chromatophores at experiment day (T_exp_) 0, 14, 28, and 42. Color denotes chromatophore age. (c) Average observation area anterior-posterior length for three squids over 6 weeks. Dashed black line is a linear fit to experimental data (5.08 + 0.24 *T*_exp_). (d) Average density and size of chromatophores in the observation area for three squids over 6 weeks. Dashed blue and orange lines are linear fit to correspondent experimental data (0.05 - 2*·*10^*−*4^*T*_exp_, and 5*·*10^*−*3^ +4*·*10^*−*6^*T*_exp_, respectively). (e) Distribution of chromatophore radii for day (*T*_exp_) 0 (*N*_tot_ = 706), 14 (*N*_tot_ = 1591), 28 (*N*_tot_ = 2715) and 42 (*N*_tot_ = 5029) from 3 squids.

### B. Aging in chromatophore growth dynamics

A stationary chromatophore size distribution over development implies that chromatophores must grow in size at a rate that depends on squid age. To prove this, we call *T*_sq_ the age of the squid, measured from the fertilization time of the egg (Fig. 6a). We also define the time *T*_exp_ elapsed from the first experimental observation, and the age 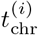 of a given chromatophore, labeled by index *i* (Fig. 6a). Given that the mantle grows linearly (Fig. 2c) and that the density of chromatophores is approximately constant during growth, the average total number of chromatophores 𝒩 (*T*_sq_) = ⟨*N*⟩ at squid age *T*_sq_ is proportional to the mantle area, which in turn grows as the square of the squid age,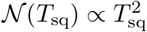. The distribution of chromatophore ages for *T*_sq_ *»* 1 is propor-tional to 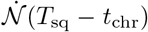, i.e., how many chromatophores were inserted at time (*T*_sq_ − *t*_chr_). After normalization, we conclude that the distribution of chromatophore ages, for large *T*_sq_, is expressed by 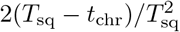.

**FIG. 6.**
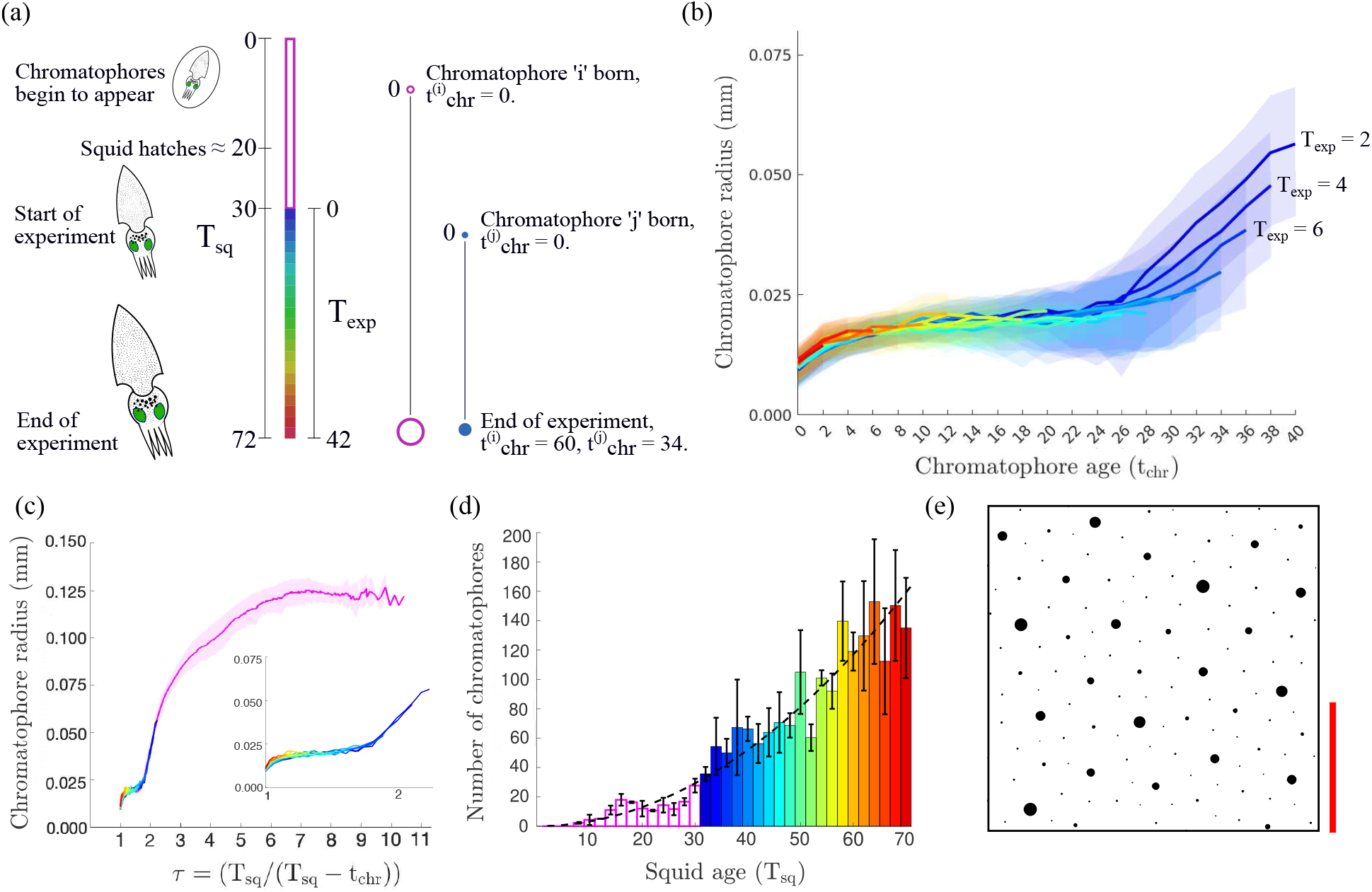
Chromatophore growth rate decays as the squid ages. (a) Timeline of squid development, where *T*_sq_ is the time measured from the fertilization of the squid egg, *T*_exp_ the time measured from the start of the experiment, and 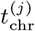 the age of chromatophore *j*. (b) Average chromatophore radius as a function of chromatophore age. Color denotes the experimental time, *T*_exp_, of chromatophore insertion (*N*_tot_ = 5733 from 3 squids followed from approximately 2-weeks-old to 8-weeks-old). (c) Average evolution of chromatophore radii as a function of chromatophore age with pre-experiment chromatophores included (*N*_tot_ = 6183 from 3 squids). (inset) Chromatophore radii from (b) plotted as function of *τ*. Inset axes are same as (c). Insertion times of all chromatophores. The black dashed line represent a quadratic fit. (e) Chromatophore pattern from simulation. The simulation area displayed is 2.5mm^2^. The red scale bar indicates 1mm.

We call *p*(*R* | *t*_chr_, *T*_sq_) the probability that a chromatophore has radius *R*, given its age *t*_chr_ and the squid age *T*_sq_. If the radius is uniquely determined by these two quantities, this probability is given by a delta function. We similarly define the probability *p*(*R*) that a randomly chosen chromatophore has radius *R*. We obtain

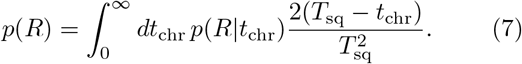

If *p*(*R*|*t*_chr_, *T*_sq_) did not depend on *T*_sq_, then Eq. (7) would predict that *p*(*R*) would not be stationary. This means that aging is necessary for achieving a stationary size distribution. If, instead, *R* is a deterministic function of *τ* = *T*_sq_*/*(*T*_sq_ *− t*_chr_), then the distribution *p*(*R*) would always be stationary. This result can be verified by directly substituting *p*(*R* | *t*_chr_) = *δ*(*R− f* (*τ*)) into Eq. (7), where *f* (*τ*) is an arbitrary increasing function. In summary, our theory predicts that the rate of chromatophore growth must slow down as the squid ages.

In agreement with our theoretical prediction, we find that, at equal chromatophore ages, chromatophores that are born earlier tend to be larger (Fig. 6b). In contrast, when plotting chromatophore radii as a function of *τ*, we obtain a precise collapse to a master curve (Fig. 6c). In this collapse, we treat *T*_sq_ as a free parameter, permitting us to estimate the value of *T*_sq_ corresponding to the start of the experiment, i.e., *T*_exp_ = 0. This value is *T*_sq_ = 30.7, which is consistent with the sum of the pre-hatching time (approximately 20 days) and the time between hatching and the beginning of our experiment (approximately 14 days). Further, by fitting individual chromatophore growth curves to the master curve, we estimate the insertion date of chromatophores that appeared before the start of our observations, see Fig. 6c and Appendix F. The complete distribution of chromatophore ages is well fitted by a quadratic law (Fig. 6d, *r*^2^ = 0.93). The predicted time for the appearance of the first chromatophore is *T*_sq_ = 6.4, which is consistent with the time of appearance of the first chromatophores on the unhatched squid after fertilization [30]. Thus, individual chromatophores reduce their growth rate as the squid ages in such a way that the distribution of chromatophore sizes is stationary during growth.

In our model, we have assumed for simplicity that the exclusion zone associated with a chromatophore is independent of its size. This assumption permitted us to study the point pattern and the chromatophore size distribution as two independent physical problems, and leads to predictions that are in quantitative agreement with our experimental observations. Despite this independence, the model is able to produce patterns in which large chromatophores tend to be surrounded by smaller ones (Fig. 6e), also in agreement with experimental observations, see Fig. 1a.

## IV. DISCUSSION

In this work, we have demonstrated how growth generates a hyperdisordered scaling of squid chromatophore patterns. By combining experimental measurements and theory, we revealed that this unconventional state of matter spontaneously arises through the interaction of random packing and growth. In particular, growth is responsible for exporting short-range disorder generated by the packing process to ever larger spatial scales. This scaling behavior is quantitatively captured by a simple system of static disks randomly placed on a growing surface, in which growth generates scale-free dynamical heterogeneity.

Given the simplicity of this mechanism, we expect that hyperdisordered behavior might be observed in other growing physical or biological systems. In contrast, it has been observed that photoreceptors form a pattern on the chicken retina which possesses hyperuniform, rather than hyperdisordered, scaling properties [19]. It has been hypothesized that this behavior may provide optimal retinal coverage properties for vision. This suggests that, depending on the interaction of cell dynamics and tissue growth, biological point patterns can be funneled into different kinds of scaling behavior.

It is natural to speculate whether hyperdisordered scaling serves a biological purpose. One hypothesis is that this type of patterning facilitates the camouflage or communication functions of the squid skin display system. Indeed, it has been shown that systems displaying fluctuations on a broad range of scales are particularly efficient at processing complex environmental signals [31]. Further, recent work on plant morphogenesis has suggested that the exporting of fluctuations to large scales via growth may increase developmental robustness [32]. Alternatively, hyperdisordered scaling might be an evolutionary spandrel [33], i.e. a side effect of the interaction of growth and packing.

Additionally, we have found that the distribution of chromatophore sizes is maintained over time by slowing down the rate of chromatophore growth as the squid ages. This result implies that chromatophores must possess some notion of squid age during growth. How this knowledge is acquired remains unclear. One possibility is that chromatophore growth rates are controlled by a morphogen that scales with system size [34–36]. Our conclusion that chromatophore dynamics must present aging is based on simple dimensional arguments. It therefore applies generally to dynamical processes on linearly growing surfaces.

In summary, our work illustrates that the interplay of dynamics and tissue growth can unlock patterns that would be impossible in non-growing tissues. Given the ubiquity of this interplay, we expect this idea to extend far beyond our particular model system.

## Supporting information

movie of a simulation of the squid model

## ACKNOWLEDGMENTS

We are grateful to the OIST Cephalopod Research Support Team for animal care, for the help and support provided by the Scientific Computing and Data Analysis and Engineering sections, Research Support Division at OIST. We thank all Pigolotti and Reiter lab members for assistance and discussion. We thank Mahesh Bandi, Joshua Shaevitz, and Salvatore Torquato for feedback on a preliminary version of this manuscript.

## Appendix A Experimental system, imaging, and chromatophore tracking

We here briefly describe our experimental model system and the imaging process. Oval squids (*Sepioteuthis lessoniana*) were bred in the OIST Marine Science Station. Juvenile squids used for this study were group housed in 50×40×40cm tanks connected to the ocean through an open seawater circulation system. The squids grow rapidly, hatching with a mantle length of approximately 16mm and reaching 90mm within 3 months. All research and animal care procedures were carried out in accordance with institutional guidelines, approved by the OIST Animal Care and Use Committee under approval number 2019-244-6.

For the analysis presented in Fig. 1, 10 animals (8 weeks post-hatching) were anesthetized (1 percent ethanol [37]), and individually transferred to a filming chamber for approximately 1 minute (Fig. 2a). They were photographed using a high resolution camera (8688 *×* 5792 pixels, approximate pixel size 7*µ*m *×* 7*µ*m). After imaging, animals were transferred to filtered seawater, and returned to their home tank after waking from anesthesia.

For chromatophore tracking, 12 animals (2-3 weeks post-hatching) were photographed in three dimensions using a 3-camera rig (Fig. 2a). Image acquisition was synchronized via an Arduino. Cameras were calibrated using a checkerboard (square size = 5mm *×* 5mm), and the reprojection error was typically less than a pixel. Images were taken every 2 days for 42 days. Squid identity was maintained across days using the spatial arrangement of chromatophores.

Within an image, chromatophore locations were determined through segmenting chromatophores using a U-net [38, 39], followed by image binarization and centroid detection. These locations were manually curated via visual inspection.

For multi-camera imaging, points corresponding to chromatophore centroids were matched through nonlinear warping of pairs of images using a custom GUI [38], followed by nearest-neighbour association of points across the two images. With chromatophore centroids identified across simultaneously acquired images, the threedimensional location of chromatophores within a day was determined using the camera calibration. Chromatophores were tracked across days in a similar manner, using nonlinear warping and nearest-neighbor matching between images separated by 2 and 4 days. A chromatophore was labelled as new if it did not have a corresponding chromatophore from the previous 2 and 4 day images.

## Appendix B Uniformity of the spatial distribution of chromatophores

We divided equally sized images of chromatophores into 16 equally sized sections and we compared the number of chromatophores in each box to both the expected number of chromatophores within an image, and across all images (total number of images = 10), using a *χ*^2^ test. The null hypothesis, that chromatophores were present in equal proportions across sections, was not rejected in all cases at significance level *β* = 0.05. We conclude that the spatial distribution of chromatophores is uniform, within our experimental uncertainty.

## Appendix C Squid model

Here, we provide details on the squid model. Chromatophore insertion is attempted by drawing new coordinate pairs (*x*_new_, *y*_new_) uniformly and at random, from within the current domain. Chromatophore insertion is successful if

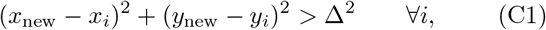

where (*x*_*i*_, *y*_*i*_) are the coordinates of an existing chromatophore *i* and the parameter Δ is the exclusion distance, i.e., the minimum distance between chromatophore centers. Otherwise, the attempt is discarded. During the simulation, chromatophore insertions are repeatedly attempted, until a maximum number of sequential failures is reached. Once this occurs, the domain length in both the *x* and *y* direction is increased by *P*_*d*_ *dt*, where *P*_*d*_ is the linear growth rate and *dt* is the simulation time step. In this growth step, the coordinates of all chromatophores are rescaled proportionally to the new domain size. Chromatophore insertion is again attempted until the maximum number of sequential failures is reached. The maximum number of sequential failures *f* is increased quadratically in time throughout the simulation, *f* = 5(6.4 + *t*^2^), to maintain a fixed number of failed attempt per unit area.

As for the parameters, the exclusion distance is set to Δ = 0.25 mm, chosen in order to match the experimental chromatophore density (see Fig. 5d). The initial domain area is 1.5Δ *×* 1.5Δ. The domain linear growth rate is *P*_*d*_ = 0.24 mm per day (see Fig. 5c). The initial simulation time is *t* = *T*_start_ = 6.4 days, in accordance with the time of first appearance of chromatophores inferred from experimental data (see Fig. 6). We set a timestep equal to *dt* = 10^*−*3^days. All simulations were written in C++ and run on the supercomputer Deigo at OIST.

## Appendix D Iterative model

Here we derive the solution of the iterative model, Eq. (4), and discuss why this expression implies that the model always presents hyperdisordered scaling. We start from with the structure factor of the initial condition

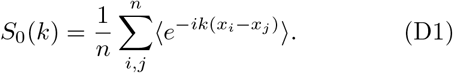

After the first iteration, we obtain

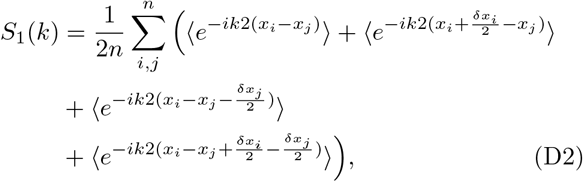

where *δ*_*x*_ are the i.i.d Gaussian random variables corresponding to the newly inserted points. We evaluate the expectations in Eq. (D2) using standard properties of Gaussian integrals, obtaining

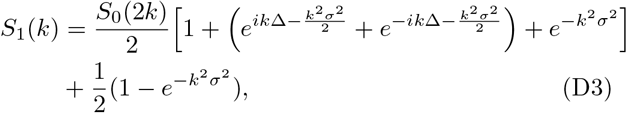

which can be simplified to

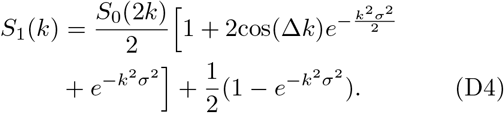

By iterating this procedure, we find that the structure factor at the *t*-th iteration is expressed by Eq. (4).

We now want to show that the structure factor diverges for *k→* 0 for an infinite system (*t→ ∞*). We set *k* = (2*π/*(*n*2^*t*^)) in Eq. (4), where *n*2^*t*^ is the number of cells after *t* iterations. We then obtain

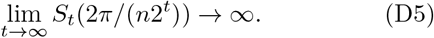

This divergence implies a hyperdisordered behavior. The mathematical reason for this divergence is that, for some *i < t*, we have that

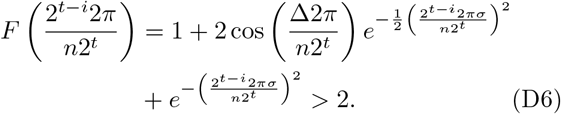

## Appendix E Kolmogorov-Smirnov test for comparing chromatophore radius distribution through time

We pooled chromatophore radii from 3 squids and binned into equally-sized bins (0, 0.01, 0.02, …, 0.18) for each *T*_exp_ *∈* (0, 2, 4, …, 42). We then performed two separate analyses using these distributions: (1) each *T*_exp_ was compared to the distribution made by aggregating chromatophore radii data across all times, (2) each *T*_exp_ was compared individually against all other *T*_exp_. In the case of (1), from 22 tests the null hypothesis was rejected 3 times at *β* = 0.05. In the case of (2), from 231 tests the null hypothesis was rejected 27 times, also at *β* = 0.05. This means that, for both analyses the null hypothesis was not rejected in approximately 85% of cases, supporting that the chromatophore size distribution is stationary.

## Appendix F Inference of chromatophore insertion times

We pooled new chromatophores for each *T*_exp_*∈* (0, 2, 4, …, 42) across 3 squids to generate the chromatophore trajectories in Fig. 6b. *T*_sq_ at *T*_exp_ = 0 was determined by minimising the residuals between chromatophore trajectories according to the function *T*_sq_*/*(*T*_sq_*− t*_chr_). This resulted in *T*_sq_ = 30.7 days. The corresponding trajectories are displayed in Fig. 6c. From these trajectories, we generated a master curve using MATLAB’s curve-fitting tool box (see below for function) and minimized the residuals of remaining chromatophores whose size coincided with this master curve using an interpolation scheme [40]. Following this procedure, we generated a new master curve, and minimized the residuals of remaining chromatophores whose size coincided with this master curve. We iterated this process until all remaining chromatophores had been matched. The final master curve is displayed in Fig. 6c.

We fitted the function describing the growth of the chromatophore radius with respect to *τ* to a 5^*th*^ degree rational using MATLAB’s curve-fitting toolbox. The equation is:

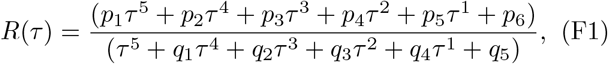

and the initial size of a chromatophore *R*(1) = *c*_*r*_ = 1.25 *·*10^*−*2^. The coefficient values are given in Table I. The support for this function is *τ ∈* [1, 7.55]. For *τ >* 7.55, we implement a linear function

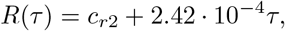

where *c*_*r*2_ = *R*(7.55) = 0.126.

**TABLE I.** Coefficient values.

|  |  |
| --- | --- |
| $p_1$ | 0.15 |
| $p_2$ | -2.21 |
| $p_3$ | 11.95 |
| $p_4$ | -27.98 |
| $p_5$ | 28.77 |
| $p_6$ | -10.25 |
| $q_1$ | -13.09 |
| $q_2$ | 56.39 |
| $q_3$ | -67.27 |
| $q_4$ | -50.63 |
| $q_5$ | 108.4 |

## Appendix G Data and code availability

Data are available from the corresponding authors on request.

